# Long-range allosteric communication, double mutant cycles, and energetic coupling in SARS-CoV-2 spike protein

**DOI:** 10.64898/2026.04.20.719543

**Authors:** Alexandra Lucas, Vamseedhar Rayaprolu, Krishna M.G. Mallela

## Abstract

SARS-CoV-2 spike protein is continuously evolving, leading to new variants. While mutations in the receptor-binding domain (RBD) enhance binding to the ACE2 receptor and evade neutralizing antibodies, the function of mutations in the N-terminal domain (NTD) remains poorly understood. Using two independent methods, surface plasmon resonance (SPR) and cryo-EM, we show that NTD mutations increase the population of spike protein with the RBD in the “up” conformational state. SPR association and dissociation kinetics of spike binding to ACE2 and antibodies, analyzed using a coupled equilibrium model, determined the relative populations and indicated that the RBD-up-to-down transition is rate-limiting relative to the RBD-down-to-up transition. Advanced model fitting of cryo-EM Coulomb potential maps confirmed the populations. The combined effect of NTD and RBD mutations exceeds the sum of their individual effects, indicating long-range allosteric communication and energetic coupling within the spike protein.

---

Coronaviruses are among the most common causes of respiratory tract infections in humans.^1,2^ In addition to exhibiting distinct cell and tissue tropisms, coronaviruses present with a wide array of clinical outcomes, ranging from the common-cold-causing HCov-OC43 to the highly fatal MERS-CoV.^1,2^ The 2002 and 2012 epidemics of SARS-CoV and MERS-CoV were notable for their increased transmissibility and mortality, but were ultimately ended by implementing public health measures and containment strategies.^3,4^ Unlike its predecessors, the 2019 SARS-CoV-2 virus has reached global pandemic status, making it the deadliest coronavirus ever to emerge. To date, SARS-CoV-2 is responsible for 7.1 million deaths worldwide and more than 780 million cases.^5^ ^6^ One factor that has enabled SARS-CoV-2 to persist, despite vaccine development,^7–10^ is its high mutational rate (1 - 2 x 10^-6^ mutations per nucleotide per infection cycle), which enables the continuous evolution of mutations that escape immunity, increase infectivity, and improve transmissibility.^8,10,11^ While the pandemic phase has ended, SARS-CoV-2 continues to evolve, leading to new variants, such as the recent nimbus and cicada variants circulating as of March 2026.

As with all coronaviruses, SARS-CoV-2 infects using surface-expressed spike glycoproteins (Fig. 1A). The SARS-CoV-2 spike binds to human angiotensin-converting enzyme II (ACE2) receptors to facilitate cellular invasion. Each spike is a homotrimer, with each monomer containing a receptor-interacting subunit and a fusogenic subunit, termed “S1” and “S2”. A specific region of each S1 subunit, called the receptor binding domain (RBD), is critical for ACE2 recognition and thus a primary target of the humoral immune response and many disease-fighting therapies.^12–15^ Each monomer in the spike trimer can exist mainly in two conformational states, with the RBD either in the “down” position or “up” position. In the “up” conformational state, RBD exposes key residues that bind ACE2 (Fig. 1A).^16^ In the “down” conformational state, the RBD binding interface with ACE2 is concealed, inhibiting virus-receptor interactions and also masking critical binding epitopes from immune targeting.^12,17^

**Figure 1.**
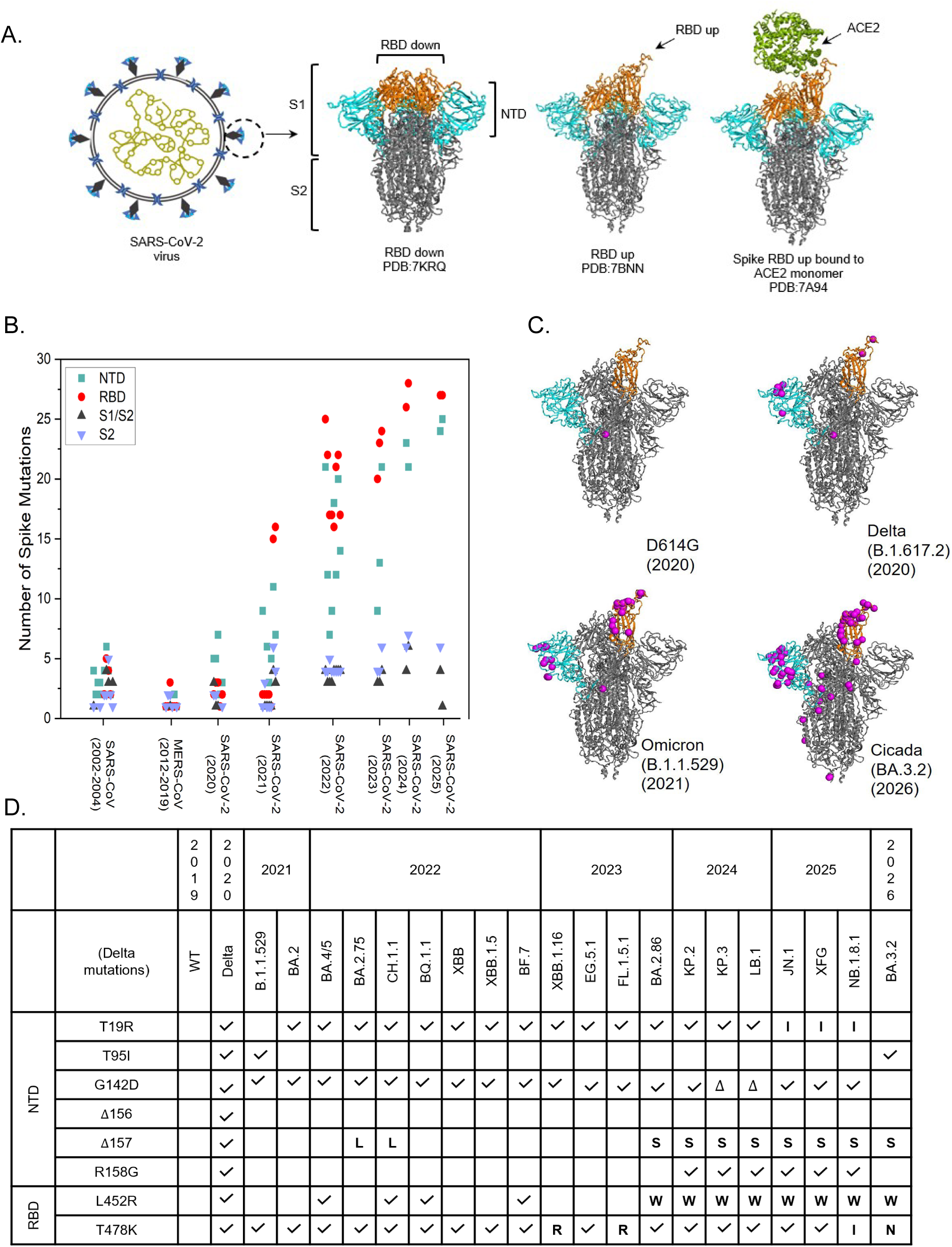
Evolution of spike protein mutations. (A) The spike protein, which forms the corona of SARS-Cov-2, exists in two conformations with RBD either in the down position or in the up position. The RBD-up conformation binds to ACE2. (B) Increasing evolution of mutations, by spike domain, across SARS, MERS, and SARS-CoV-2 viruses. (C) Increasing evolution of mutations across SARS-CoV-2 variants. Mutations are represented as spheres in one monomer in the spike trimer for clarity and show how mutations are generally localized to specific regions on each domain. (D) Table showing the prevalence of the Delta NTD and RBD mutations across multiple variants that have evolved from 2020 to 2026. Some variant mutations occurred at the same position as in Delta, but with a different amino acid change. These are shown using the amino acid letter identifier.

Because the spike protein is the primary mechanism by which SARS-CoV-2 invades human cells, it has consistently evolved function-enhancing mutations across different coronavirus strains. The most frequently mutated region is the RBD (Fig. 1B), as mutations in this domain can directly enhance spike protein binding to ACE2 or enable escape from neutralizing antibodies targeting RBD epitopes.^18^ The N-terminal domain (NTD) (Fig. 1A) is the second most frequently mutated region in the spike protein (Fig. 1B). NTD does not directly interact with ACE2. However, NTD has evolved an increasingly diverse constellation of mutations over the course of the pandemic, which are distal to the RBD mutations (Fig. 1C).^15,19–21^ NTD’s extensive mutational capacity may contribute to viral fitness, as both SARS-CoV and MERS-CoV have also evolved NTD mutations.^22–25^ With each new SARS-CoV-2 variant, the number of NTD mutations has increased, similar to that in the RBD (Figs. 1B and C). Many of the NTD mutations are present across multiple spike protein variants with increased infectivity and transmissibility (Fig. 1D).^26^

Initial research suggested that NTD mutations evolved solely to promote immune evasion.^27,28^ However, this hypothesis does not adequately explain all NTD mutations. Alternative theories have proposed that NTD and its mutations facilitate viral-cell fusion in a manner similar to that of other coronaviruses. However, no research has conclusively proven this, and many published studies report conflicting results.^29^ One recent study showed increased virus-cell entry and spike cleavage with NTD mutations.^30^ Another study demonstrated that swapping the distal NTD loops of SARS-CoV and SARS-CoV-2 altered cellular entry of pseudoviruses. These findings are especially intriguing given that many NTD mutations are in the loops that are distal to RBD (Fig. 1C). While these works provide novel insights into how alterations in NTD may impact cellular infection, no experimental work has specifically studied the impact of NTD mutations on molecular mechanisms by which the spike protein interacts with ACE2 and neutralizing antibodies, though some *in silico* studies suggested long-range communication between NTD and RBD.^31,32^

To address how NTD mutations affect the spike protein’s structure and function, we constructed four chimeric proteins with varying NTD and RBD domains starting from the wild-type and Delta variant spike proteins (Fig. 2A). We chose the Delta variant because of the appearance of novel mutations that were not seen in earlier variants (Alpha, Beta, and Gamma), and these mutations have continued to appear in subsequent Omicron and its sub-lineages that have been circulating (Fig. 1D). In addition, Delta was considered the most dangerous variant of SARS-CoV-2, because it combined very high transmissibility, twice that of the original Wuhan strain, and about as infectious as chickenpox, a much higher viral load (up to ∼1000-fold), and increased severity of disease with a higher risk of hospitalization. Chimeric spikes were constructed with WT S2 subunit and either the wild-type or Delta variant RBD or NTD (Fig. 2A), enabling analysis of the individual and combined domain mutations. We examined the effect of individual domains on spike protein expression, stability, and binding to the ACE2 receptor and to the three FDA-approved therapeutic neutralizing antibodies. We used two independent methods, surface plasmon resonance (SPR) and cryo-EM, to reach the same conclusions about how NTD affects RBD conformational dynamics. Analyzing the SPR association and dissociation kinetics using a coupled equilibrium model, rather than the conventional 1:1 binding model used in most published studies, allowed us to determine the relative populations of the different conformational states of the spike protein, which were confirmed by advanced model fitting of cryo-EM Coulomb potential maps.

**Figure 2.**
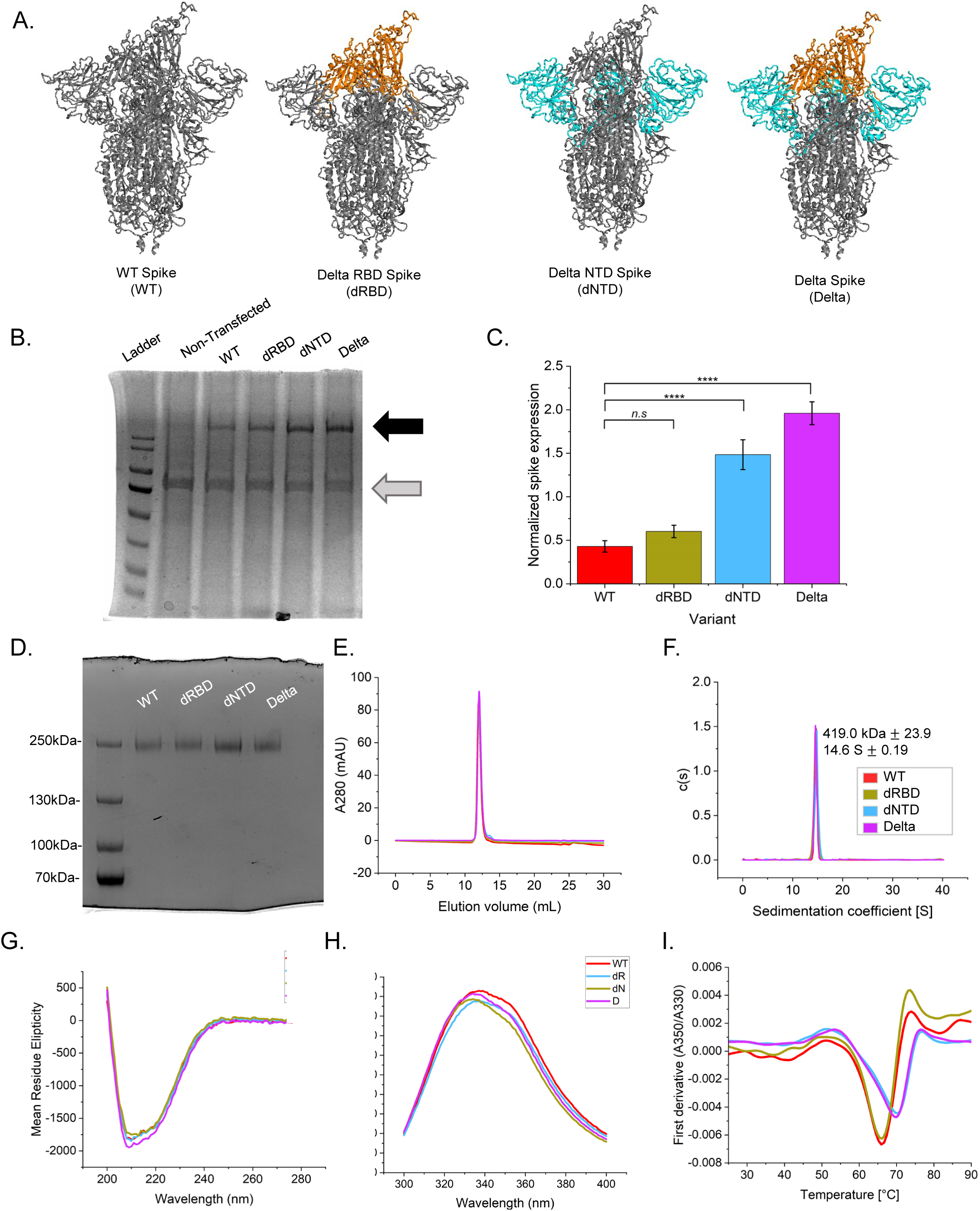
Characterization of WT and mutant spike proteins. (A) The four chimeric spikes used in this study: WT with wild-type RBD and NTD domains, dRBD with wild-type NTD and Delta RBD, dNTD with Delta NTD and wild-type RBD, and Delta with both Delta NTD and RBDs. (B) SDS-PAGE showing the relative expression of the four chimeric proteins (black arrow). A non-transfected culture was used as a negative control. Ladder represents molecular weight markers of 180, 130, 100, 70, 55, 40, 35, and 25 kDa (from top to bottom). (C) Relative expression of each spike protein was normalized to a natively expressed host protein (gray arrow) and plotted. Results in panel B represent the average of 5 replicates for each protein. Statistical significance was determined via a two-sample T-test. Significance is represented as follows: *p<0.05 (*), p<0.01 (**), p<0.001 (***), p<0.0001 (****).* (D) SDS-PAGE gel and (E) size exclusion chromatograph (SEC) showing the homogeneity of purified spikes. (F) Sedimentation velocity distribution plots from analytical ultracentrifugation show that all four chimeric proteins are trimers. (G) Far-UV circular dichroism spectra and (H) intrinsic aromatic fluorescence spectra show no effect of mutations on global protein structure. (I) Differential scanning fluorimetry (DSF) thermal melt showing changes in the fluorescence ratio at emission wavelengths of 350 nm and 330 nm as a function of temperature. The maxima and minima represent inflection temperatures listed in Table 1.

**Table 1:** T_m_ values from DSF thermal melts (Fig. 2I)

| Variant | $T_m$ #1 | $T_m$ #2 | $T_m$ #3 |
| --- | --- | --- | --- |
| WT | $50.94 \pm 0.78$ | $65.95 \pm 0.11$ | $74.14 \pm 0.14$ |
| dRBD | $50.63 \pm 1.25$ | $65.83 \pm 0.11$ | $73.76 \pm 0.14$ |
| dNTD | $52.44 \pm 0.70$ | $69.95 \pm 0.22$ | $77.06 \pm 0.12$ |
| Delta | $53.17 \pm 0.53$ | $69.59 \pm 0.04$ | $76.91 \pm 0.41$ |

## Results

### NTD mutations increase spike protein expression

Improved expression of viral proteins confers a selective advantage by enabling the production of more viral particles, potentially enhancing infection.^33^ Clinical reports on the Delta variant showed elevated viral titers in vaccinated and unvaccinated patients.^34^ We tested the expression of four chimeric proteins (Fig. 2A): WT with the wild-type NTD and RBD domains of the original Wuhan strain; dRBD with wild-type NTD and Delta RBD; dNTD with Delta NTD and wild-type RBD; and Delta with both NTD and RBD as in the Delta variant. All four proteins were constructed on the HexaPro spike protein background, which was designed to overcome low protein expression.^35^ Expression was tested in suspension Expi293 HEK cells and quantified by SDS-PAGE analysis^36^ (Fig. 2B). Both chimeric proteins, dNTD and Delta, containing the Delta NTD, showed statistically significant increases in spike protein expression, with a 4-fold and 3-fold increase compared to the WT, respectively (Fig. 2C). No significant effect on protein expression was observed for the chimeric protein dRBD that contains only the Delta RBD domain. This is consistent with previous results showing that Delta mutations enhance protein expression, although the effect of RBD mutations was comparatively lower for the full-length trimeric spike protein than for the monomeric RBD.^36^ The enhanced expression of chimeric proteins containing the Delta NTD indicates that NTD mutations contribute to increased spike expression, with the high expression yield of the Delta variant resulting from the combined effects of RBD and NTD mutations (Fig. 2C).

### NTD mutations do not affect spike structure, but do impact its stability

Spike purity was confirmed using SDS-PAGE (Fig. 2D) and analytical size exclusion chromatography (SEC) (Fig. 2E) prior to characterizing its structure and function. SDS-PAGE showed a single band, and SEC showed a single peak, showing homogeneity of the four chimeric proteins. Sedimentation velocity analytical ultracentrifugation (SV-AUC) experiments confirmed that all spikes were intact trimers (Fig. 2F) with the molecular weight estimated from the sedimentation coefficient matching that of the calculated molecular weight of the spike trimer. Circular dichroism (Fig. 2G) and intrinsic fluorescence (Fig. 2H) analysis showed that mutations did not significantly affect the secondary and tertiary structures of the spike protein, respectively.

Differential Scanning Fluorimetry (DSF) was used to assess the effect of mutations on the spike stability. Distinct unfolding profiles were observed for all four chimeras, with melting temperature (*T_m_*) shifting towards higher temperatures in spike proteins with mutant NTDs (Fig. 2I & Table 1). RBD mutations had no detectable effect, consistent with our previous work on the Delta variant monomeric RBD.^36^ The improved stability of dNTD and Delta chimeric proteins suggests that NTD mutations increase spike protein stability, which correlates with their increased expression (Fig. 2C).

### NTD and RBD mutations enhance spike affinity for ACE2

Surface plasmon resonance (SPR) was used to determine the association and dissociation kinetics of chimeric spike proteins binding to the human ACE2 receptor. In these experiments, the spike protein was covalently attached to the SPR chip, and ACE2 was used as the analyte with varying concentrations to determine the association kinetics. For dissociation kinetics, buffer was used as the analyte. Fig. 3A shows the response curves (sensorgrams), which were initially fit using a 1:1 Langmuir model. However, the association curves deviated clearly from 1:1 binding behavior. This deviation could not be explained even after incorporating the mass transport limitation (MTL) in SPR experiments^37^ (Fig. 3B). Because spike protein is a trimer and the RBD of each monomer can, in principle, bind to one ACE2, the observed deviation from 1:1 behavior likely represents a multiphasic binding. A more complex model is required to correctly investigate spike protein binding kinetics to ACE2. We employed an analytical model determination method to accurately identify the best model for interpreting spike protein binding to ACE2.^38^ SPR experiments with 8 analyte concentrations were performed for WT spike binding to ACE2 (Fig. 3C). These sensorgrams were fit to a model-independent biphasic kinetics, and the goodness of the fit is evident from the residuals. The obtained rate constants (s1 and s2), their sum, and their product were plotted as a function of analyte concentration (Figs. 3D-G). Because the observed rate constants vary differently with analyte concentration for each kinetic scheme (Fig. 4A), these trends were used to determine the kinetic mechanism by which the spike protein binds to ACE2. The three simplistic models we tested that can result in biexponential kinetics were the two-state conformational change model where RBD transitions from down to up conformation before binding to ACE2 (Model 1), heterogeneous ligand model where the spike trimer can bind independently to two ACE2 molecules (Model 2), and a bivalent analyte model where the two ACE2 molecules bind sequentially to the spike trimer (Model 3) (Fig. 4A). These models were chosen based on the prior experimental knowledge that (1) spike trimer exists in multiple states with either RBD in the down or up confirmation (Fig. 1A)^14,39^ and (2) multiple ACE2 molecules can bind to the spike trimer with one ACE2 binding to the RBD of a monomer in spike trimer.^39,40^ Since spike is a trimer with each monomer existing either in the RBD up or down conformations, more complex kinetic schemes can be written, but we first tested whether any of these three simple models can explain the SPR kinetics.

**Figure 3.**
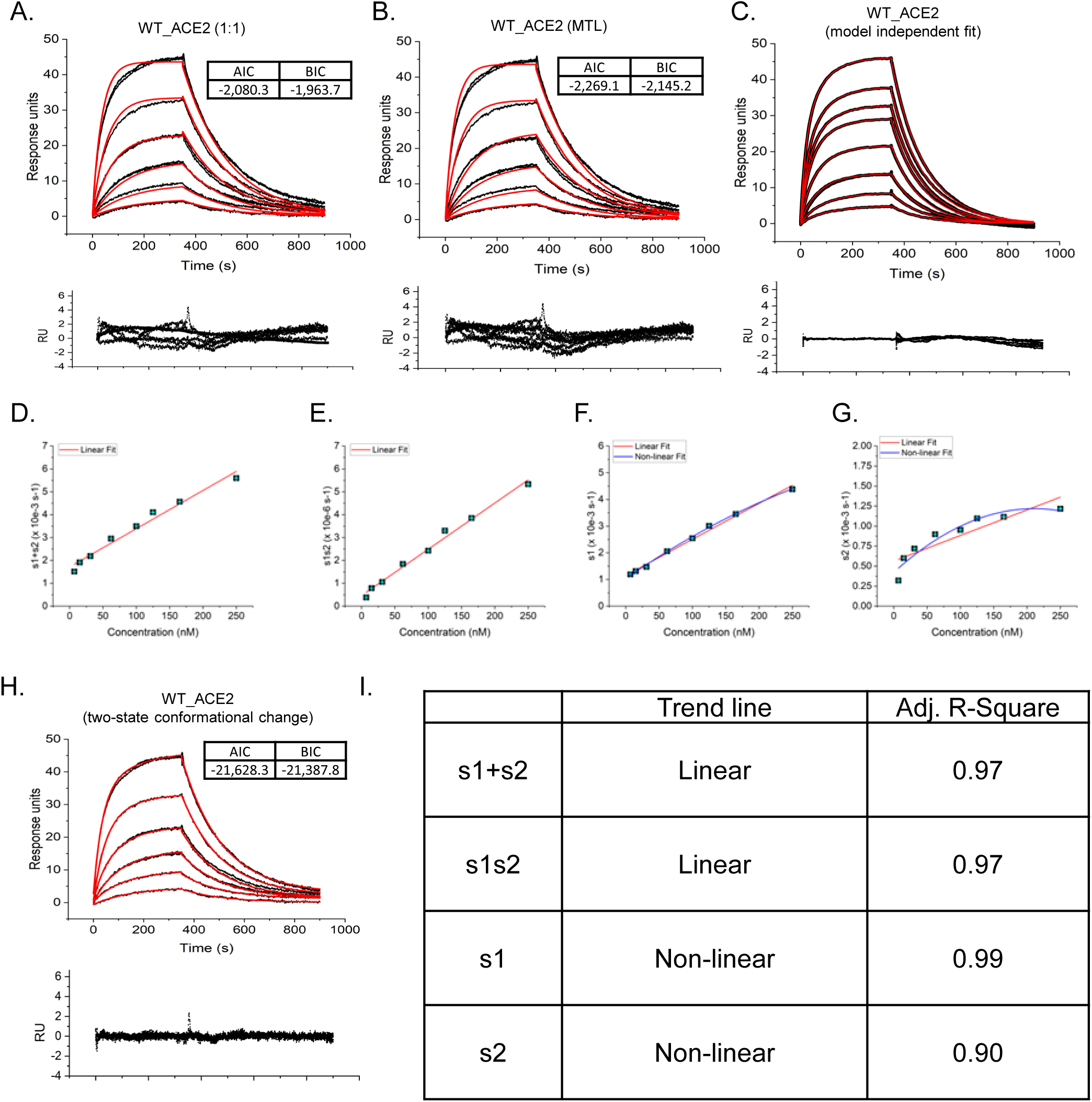
Model comparison for fitting SPR binding curves of WT spike binding ACE2. Data curves are shown in black and fit lines in red. Sensorgrams were fit with (A) a 1:1 model, and (B) a mass transport limited (MTL). The two observed rates (s1 and s2) from fitting the sensorgrams in panel C with a model-independent biexponential model, were plotted as a sum (panel D), product (panel E), and individual rate constants (panels F and G) with respect to the analyte ACE2 concentration. Visual inspection of plots D-G allowed the selection of a model from Fig. 4A. The selected model was then used to fit WT and ACE2 binding (panel H), showing good fits and random residuals. The concentrations of ACE2 from top to bottom are 250 nM, 125 nM, 62.5 nM, 31.25 nM, 15.625 nM, 7.8125 nM. The sensorgram (panel C) used for model-independent analysis included additional ACE2 concentrations of 100 nM and 165 nM. AIC and BIC values show the statistically “best fitting” model in terms of goodness of fit and model complexity, with lower values indicating the more appropriate model. Panel I shows the trend line type and adjusted R-Square values for panels D-G.

**Figure 4.**
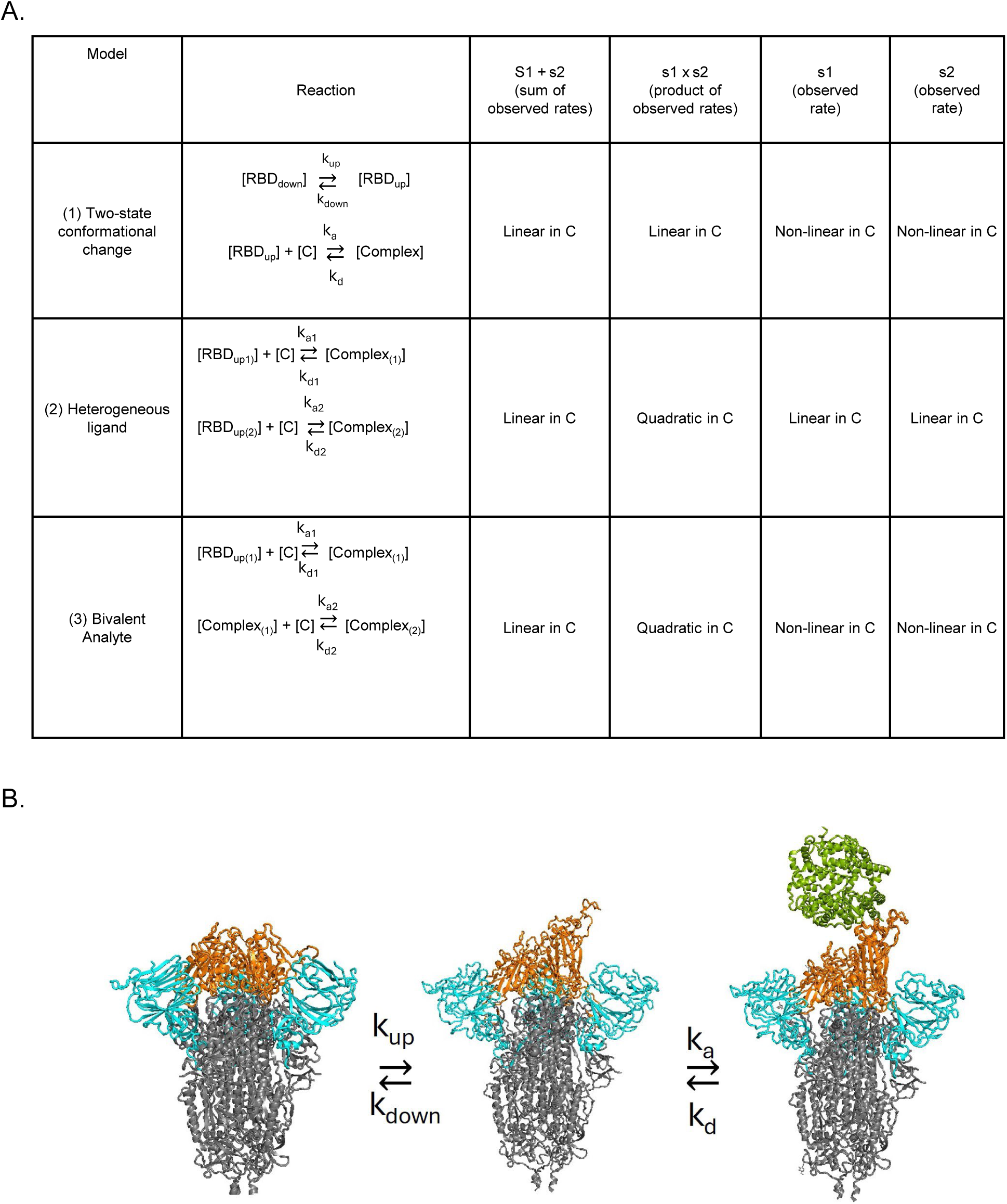
Model determination reactions and plot trends. (A) Table describing the three simplistic reaction schemes tested with WT spike sensorgrams binding to ACE2. (B) Graphical schematic of the two-state conformational change model selected for fitting the sensorgrams of spike chimeras binding to ACE2 and class 1 antibodies.

Linear regression of the observed rate plots (Figs. 3D-3G, 3I) as a function of increasing ACE2 concentration showed a clear linear trend in the sum and product of the rate constants. However, the individual rate constants showed a non-linear trend, which implies that Model 1 involving a conformational change from RBD-down to RBD-up state prior to binding to ACE2 is the appropriate model for interpreting SPR kinetics (Fig. 4B). This was further confirmed by re-fitting spike-ACE2 sensorgrams with Model 1, which showed much improved fit and residuals compared to 1:1 and MTL (Fig. 3H vs. Figs. 3A & 3B).

SPR sensorgrams for all four chimeric proteins binding to ACE2 were fit with a two-state conformational change model. The association rate constant (k_a_), dissociation rate constant (k_d_), the rate constants of RBD transitions (k_up_ and k_down_) obtained from the model fits, the calculated equilibrium constant for the conformational change (K=k_down_/k_up_), and the dissociation constant (K_D_bind_ =k_d_/k_a_) are plotted in Fig. 5. The overall binding affinity was calculated as K_D_= K*(1+ K_D_bind_). Results showed that dRBD and WT spikes have comparable ACE2 affinity (K_D_) (Fig. 5H), consistent with earlier published reports on isolated Delta RBD that do not affect ACE2 binding.^36,41^ Interestingly, dNTD showed a 1.6-fold greater affinity to WT, indicating that NTD mutations significantly affect RBD-ACE2 interactions. Delta spikes showed the strongest overall affinity, indicating that combined NTD and RBD domain mutations have a greater impact than individual domains. To explore this further, we compared the rate of RBD conformational changes (Fig. 5E). The results showed that the forward rate (k_up_) of RBD changing from down to up did not differ significantly across chimeras. However, the reverse rate (k_down_) of RBD-up transitioning to RBD-down was slower in the presence of NTD mutations. Thus, NTD mutations appeared to stabilize the RBD-up state of the spike protein, which would explain the increased affinity observed for dNTD and Delta chimeras.

**Figure 5.**
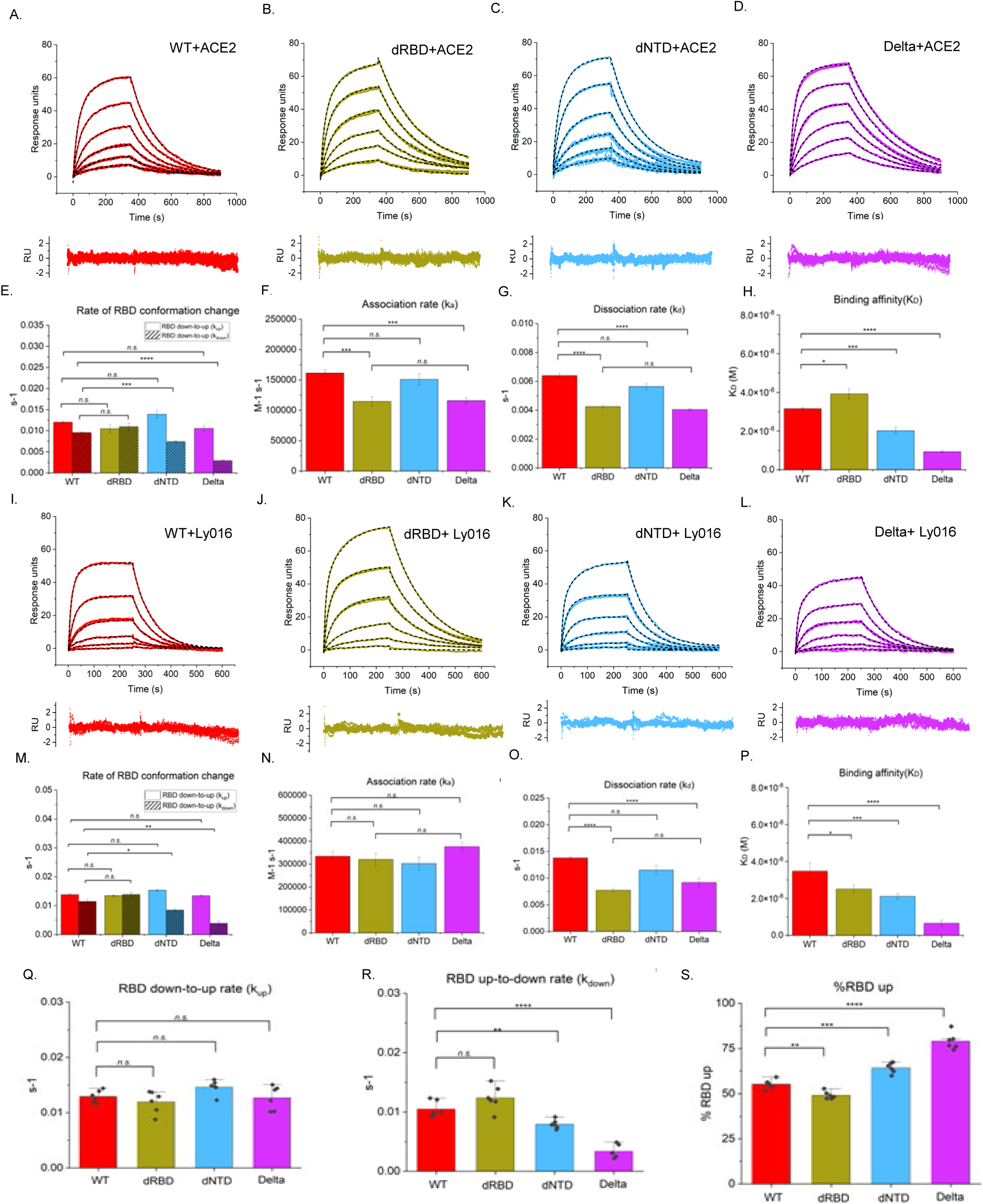
Spike chimeras binding to ACE2 and the ScFv of a class 1 antibody Ly016. Sensorgrams of WT (red), dRBD (green), dNTD (blue), and Delta (pink) spike chimera binding ACE2 (A-D) or Ly016 (I-L), respectively. ACE2 concentrations from top to bottom were 250nM, 125nM, 62.5nM, 31.25nM, 15.625nM, 7.8125nM.Ly016 concentration from top to bottom are 300nM, 100nM, 33.33nM, 11.11nM, 3.7nM, 1.1234nM. Bar graphs show (E & M) the rate of RBD conformational change (k_up_, k_down_), (F & N) association rates (k_a_), (G & O) dissociation rates (k_d_), and (H & P) the complex dissociation constant (K_D_) determined from fitting for ACE2 binding (A-D) and Ly016 binding (I-L), respectively. The combined average rate constants of RBD transitioning from down-to-up (panel Q) and from up-to-down (panel R) calculated from fitting ACE2 and Ly016 binding sensorgrams. Panel S shows the calculated percentage of RBD-up conformation of the spike determined from the rates shown in panels Q and R. Dots are individual rate values from the ACE2 and Ly016 datasets, while the bars represent the average. Results represent the average of 3 replicates for each chimera. Statistical significance was determined via two-sample T-test. Significance is represented as follows; *p<0.05 (*), p<0.01 (**), p<0.001 (***), p<0.0001 (****)*.

In terms of the effect of mutations on the association and dissociation kinetics of ACE2 binding, NTD mutations did not affect the kinetics (WT → dNTD and dRBD → Delta in Figs. 5F and 5G), as expected since ACE2 binds to RBD and not to NTD. However, RBD mutations slowed both association and dissociation kinetics (WT → dRBD and dNTD → Delta).

### NTD and RBD mutations enhance spike affinity for a Class 1 antibody

Neutralizing antibodies identified in human animal models and in the serum of recovered COVID-19 patients were developed as therapeutic drugs and approved by the FDA for emergency use. These have been classified into different classes based on where they bind to the RBD.^42^ Class 1 antibodies directly compete with ACE2 for binding to the spike protein and bind predominantly to the RBD-up conformational state. We used the single-chain variable fragment (ScFv) of Eli Lilly’s Ly016 antibody to examine how the NTD and RBD mutations affect the spike protein binding to Class 1 antibodies. Because Class 1 antibodies bind to the spike protein in a similar manner to ACE2, the same two-state conformational model (Model 1 in Fig. 4A) was used to analyze binding (Fig. 5I-P). Both dRBD and dNTD showed increased affinity, and the Delta variant displayed the strongest binding.

For association kinetics, no significant differences were observed among the four chimeric proteins (Fig. 5N). However, the dissociation kinetics were significantly slowed by the presence of RBD mutations, but not the NTD mutations (Fig. 5O).

### NTD mutations increase the RBD-up conformation of the spike protein

Since the equilibrium between RBD-down and RBD-up conformational states of the spike protein should be independent of which ligand it binds to (Model 1 in Fig. 4A), the k_up_ and k_down_ rate constants should be the same for both ACE2 and Class 1 antibody binding. Figs. 5E and 5M confirm that these rate constants were similar for both ACE2 and Ly016 binding for each individual chimeric protein. The rate constants are plotted together in Figs. 5Q and 5R. The individual rate constants indicate that neither NTD nor RBD mutations significantly affected the kinetics of the spike protein conformational transition from RBD-down to RBD-up (Fig. 5Q). The major difference is in the RBD-up to RBD-down rate constant (Fig. 5R). NTD mutations make the spike protein stay in the RBD-up state for a longer time. Both rate constants were used to calculate the fraction of RBD-up in different chimeric proteins (Fig. 5S). Compared with WT, RBD mutations (dRBD) decrease the population of the RBD-up state, whereas NTD mutations (dNTD) significantly increase it. In the presence of both RBD and NTD mutations (Delta chimeric protein), the RBD-up population increase is much higher.

### NTD mutations did not affect spike binding to Class 2 and Class 3 antibodies

Class 2 and Class 3 antibodies bind to the spike protein in both RBD-down and RBD-up states.^42^ Figs. 7A-7D and Figs. 7H-7K show the SPR association and dissociation kinetics of the four chimeric proteins binding to ScFvs of a Class 2 Eli Lilly’s antibody drug Ly555 and a Class 3 Regeneron’s antibody drug REGN10987, respectively. These data were analyzed using a 4-state conformational change Model 4 (Fig. 6A), which is a modified version of Model 1 in Fig. 4A, with an additional arm accounting for antibody binding to the RBD-down state of the spike protein. The rate constants, k_up_ and k_down_, were fixed for each chimera to the average rates determined from fitting the ACE2 and Ly016 binding (Figs. 5Q & 5R), to minimize the number of fit parameters. Since the analytical solutions to this 4-state kinetic scheme are difficult to fit with conventional data-fitting software, such as OriginPro, due to the large number of parameters, we used MATLAB’s built-in compartmental analysis (Fig. 6B) to fit the data numerically.

**Figure 6.**
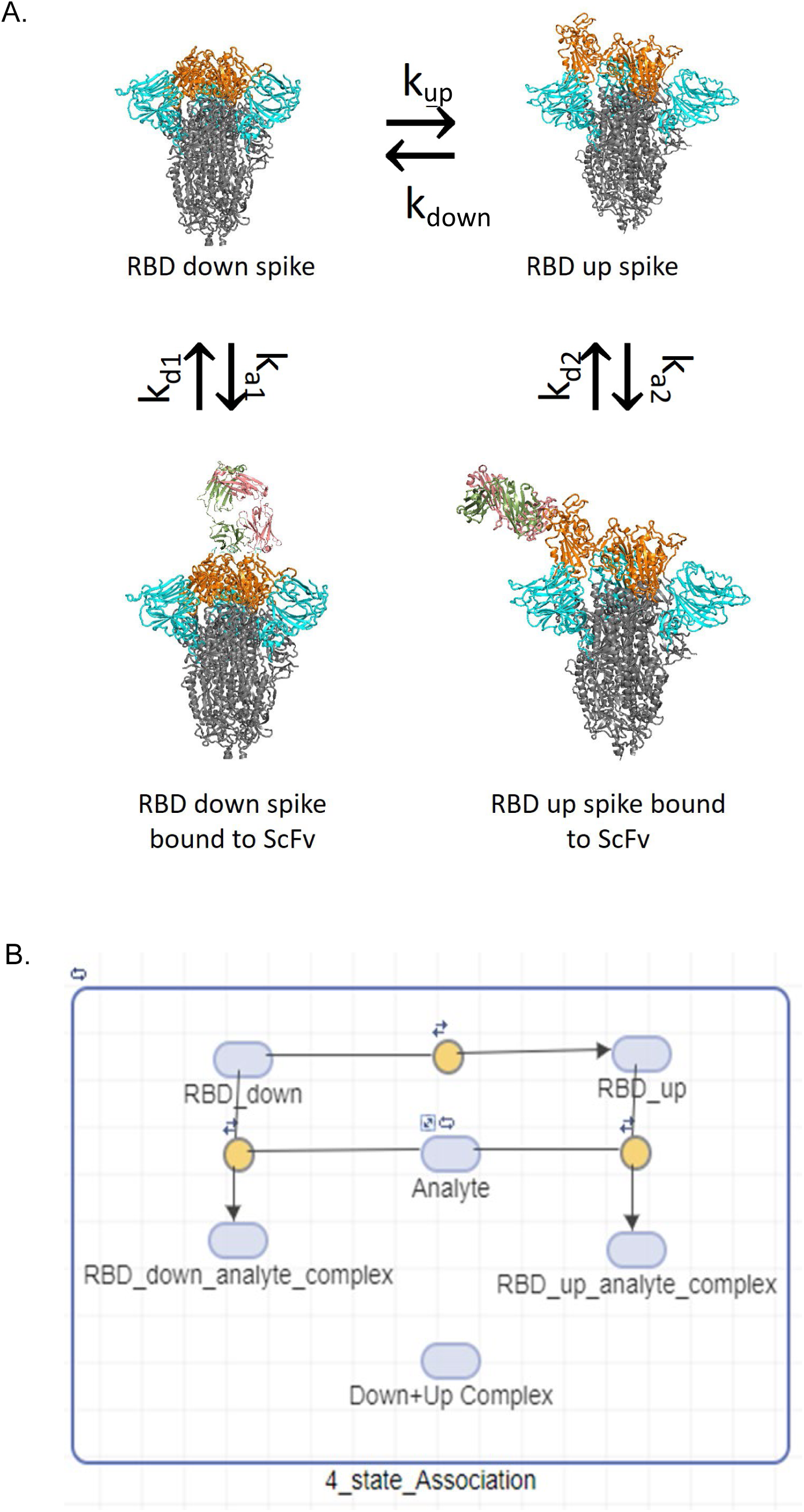
(A) 4-state kinetic scheme for spike protein binding to class 2/3 antibodies. (B) Compartmental model in MATLAB for numerically fitting the SPR data (Fig. 7) to the model in panel A. The fit lines represent the Down+Up Complex, which is the sum of the two RBD_analyte_complexes from each RBD state, specified in MATLAB by the Algebraic rule (o=Down+Up Complex - (RBD_down_analyte complex + RBD_up_analyte_complex)).

**Figure 7.**
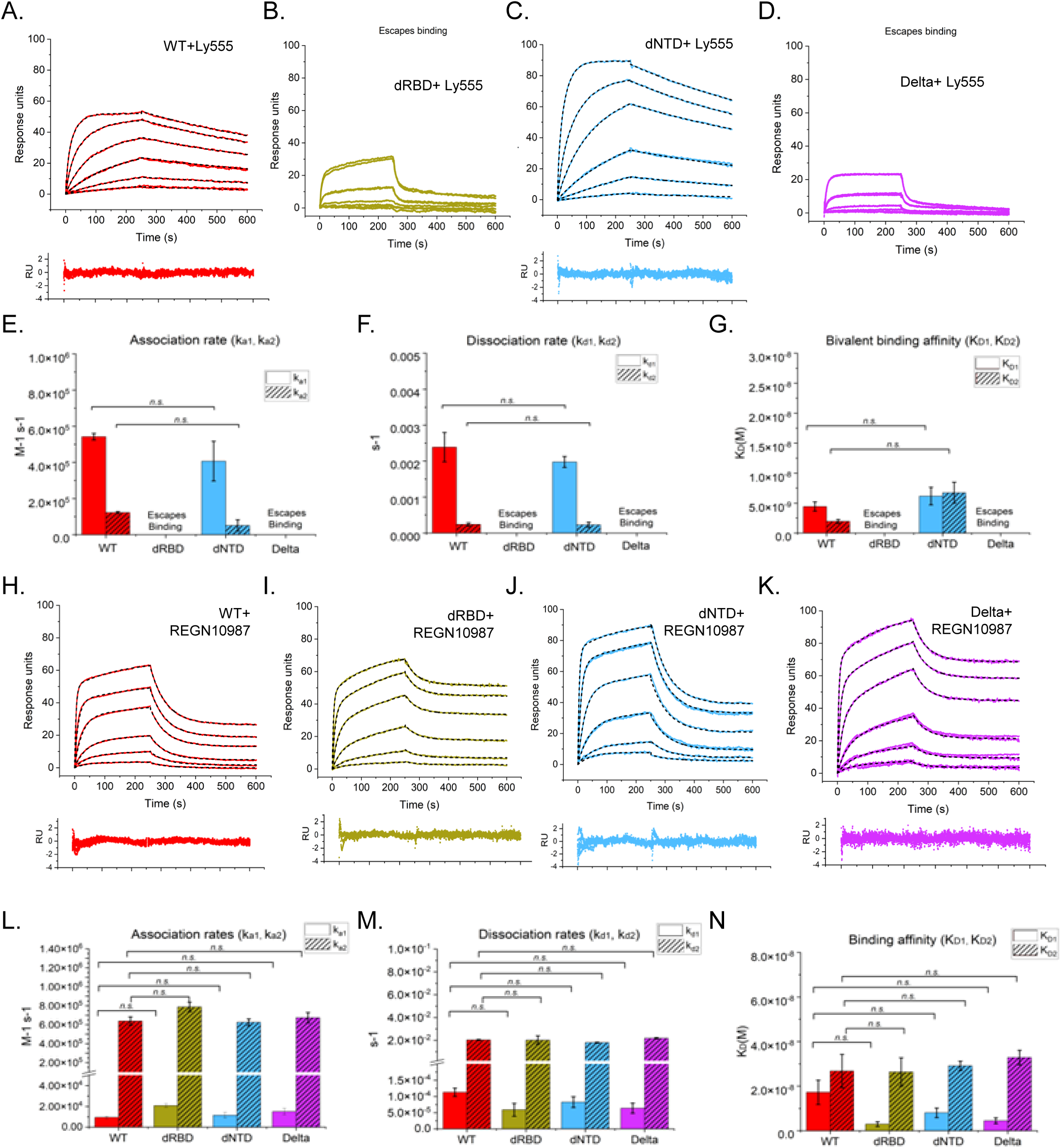
Spike chimeras binding to the ScFvs of a class 2 antibody Ly555 and a class 3 antibody REGN10987. Sensorgrams of WT (red), dRBD (green), dNTD (blue), and Delta (pink) spike chimera binding Ly555 (A-D) or REGN10987 (H-K), respectively. ScFv concentrations from top to bottom are 300nM, 100nM, 33.33nM, 11.11nM, 3.7nM, 1.1234nM. Bar graphs display the two association rates (ka1,ka2), dissociation rates (kd1,kd2), and K_D_‘s (K_D1_, K_D2_) determined from fitting the Model 4 to binding Ly555 (E-G) and REGN10987 (L-N). Binding affinity values (K_D1_ and K_D2_) represent the ratio of dissociation over association for each reaction. Statistical significance was determined via two-sample T-test. Significance is represented as follows; *p<0.05 (*), p<0.01 (**), p<0.001 (***), p<0.0001 (****)*.

Figs. 7E – 7G show the fit parameters for the four spike chimeric proteins binding to the ScFv of Class 2 Ly555 antibody. Consistent with our earlier studies on isolated RBD,^18,36^ the two chimeras containing RBD mutations, dRBD and Delta, showed no or very weak binding to the spike protein. More importantly, no significant difference was observed in the association and dissociation kinetics, or the binding affinity between the WT and dNTD chimeras, indicating that an increase in the RBD-up state population of the spike protein due to NTD mutation should, in principle, not affect the antibody binding to either the RBD-down state or the RBD-up state of the spike protein.

Figs. 7L – 7N show the association and dissociation rate constants and the equilibrium dissociation constants for the four spike proteins binding to the Class 3 REGN10987 ScFv. No statistically significant differences were observed between WT and dNTD, indicating that NTD mutations do not affect spike protein binding, consistent with the results for Class 2 ScFv. In contrast to Class 2 Ly555 ScFv, RBD mutations did not affect the spike protein’s binding to Class 3 ScFv.

### Cryo-EM structures confirm the conclusions from SPR sensorgrams

The main conclusion from the above SPR analysis of binding of the four chimeric spike proteins to ACE2 and three neutralizing antibodies is that the NTD mutations increase the population of RBD-up states of the spike protein compared to the RBD-down states. We used cryo-EM structural analysis to validate the calculated percentages of RBD-up and RBD-down states of the spike protein from SPR experiments. With RBD and NTD the most flexible regions, particularly in the RBD-up conformations of the spike protein and the presence of mutations mainly in the loop regions, determining the relative populations of RBD-down and RBD-up states is not trivial (Fig. 8). First, motion correction and contrast transfer function (CTF) estimation were performed, followed by micrograph curation to remove low-quality micrographs. Several rounds of particle picking and 2D classification were performed to isolate classes with clear spike structures, which were then used for ab initio 3D reconstruction. Reconstructed particles were subjected to heterogeneous refinement, which revealed an all-RBD down state and a 1-RBD up state. Each state was separately C3-symmetry expanded, and 3D classification with a focused mask over the RBD was used to separate 1-RBD-up particles further. The process was repeated with each subsequent 3D class to separate all 1-up from all-down states, until the final classes were virtually homogeneous. All final classes corresponding to each RBD state underwent a final non-uniform refinement (1-RBD up particles) or homogeneous refinement with C3 symmetry (all-RBD down particles). In cases where volumes showed missing resolution in RBD, a local refinement was run with a focused mask over RBD, with the fulcrum coordinates set at the base of the mask. Finally, local resolution estimation was run on all final volumes. The structures were determined at a global resolution between 2.3 and 4.0 Å across all chimeras. The resolution was not homogeneous, ranging from 2 Å in the stable S2 core to 6 Å in the RBD and NTD.

**Figure 8.**
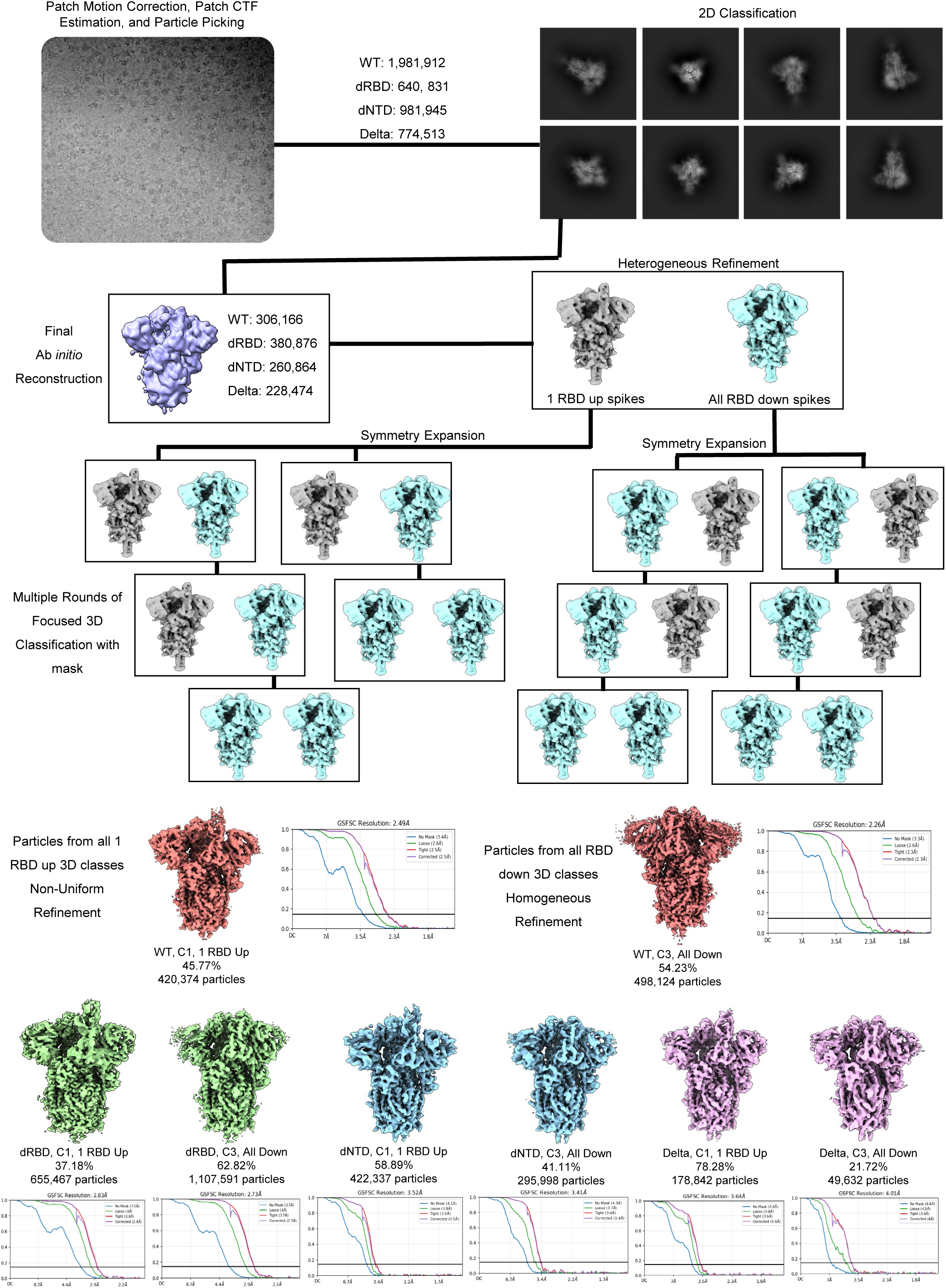
Cryo-EM workflow schematic of determining the relative populations of RBD-down and RBD-up conformations of the spike protein. EM maps were motion corrected and CTF estimated using CryoSparcv4. 2D classification separated particles for 3D construction. Populations were determined from 3D ab-initio constructed maps, and iterative 3D classification using a focused mask over RBD. Final maps produced using Local Refinement.

Fig. 9A shows the cryo-EM structures of the RBD-up and RBD-down conformations of the four spike chimeric proteins. The Coulomb potential maps captured either an all-RBD-down conformation or a 1-RBD-up conformation. No 2- or 3-RBD-up conformations were observed in our data, consistent with the SPR results (Figs. 4 & 5). For WT, ∼46% of particles corresponded to the 1-RBD up state, and 54% were in the all-down state. Focused refinement and 3D variability performed on the RBD-up particles enhanced the resolution, and we could visualize the range of RBD motion. The RBD conformation was observed to be very flexible, as is common with spike proteins. Similar analysis of Delta showed 78% in the 1-RBD-up state, consistent with SPR results indicating that Delta has a larger RBD-up population than WT. For the dNTD and dRBD chimeras, 59% and 37% of particles were observed in the up conformation. Thus, the spikes with NTD mutations showed an increase in the RBD-up population, whereas RBD mutations showed a decrease in the RBD-up population. Furthermore, the populations of RBD-up states were comparable and followed similar trends to those calculated from SPR experiments (Fig. 5).

**Figure 9.**
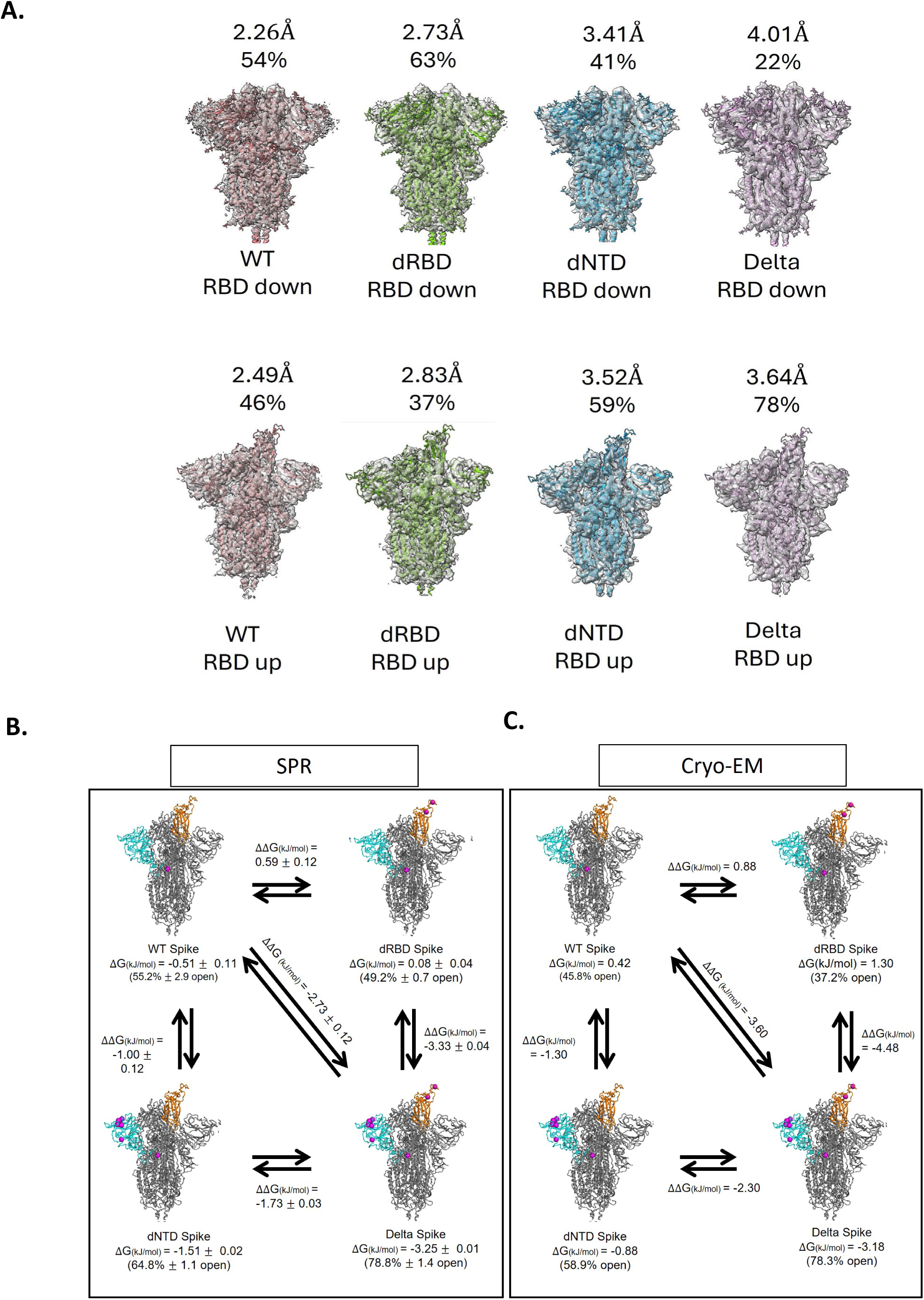
(A) RBD-down (top panels) and RBD-up (bottom panels) conformational states of the four chimeric spike proteins determined by cryo-EM. (B) Double mutant cycles showing shifts in conformational free energy calculated from SPR sensorgram fits and Cryo-EM populations. The combined effect of NTD and RBD mutations (WT → Delta) is higher than that of the sum of free energy changes due to NTD mutations only (WT → dNTD) and RBD mutations only (WT → dRBD).

### Long-range allosteric communication and energetic coupling between NTD and RBD mutations

NTD and RBD mutations are at distal sites in the spike protein (Fig. 1C). Even though the mutated residues are not in contact with one another, both SPR (Fig. 5) and cryo-EM (Fig. 9) methods show that NTD mutations increase the population of RBD-up conformation of the spike protein, implying long-range allosteric communication between these two domains. To quantify the thermodynamics of such allostery, we used conventional double mutant cycles.^43^ Figs. 9B & 9C show the double mutant cycles constructed from SPR and cryo-EM determined populations, respectively. Both techniques work on different principles. In SPR, the spike protein was immobilized on the chip using covalent chemistry, the analyte was flowed at varying concentrations, and the change in mass due to binding was detected in response units. Cryo-EM uses flash-freezing of samples to visualize the trapped conformational states via Coulomb potential maps. Hence, we do not expect identical populations of the up and down states between the two techniques; rather, we look specifically for trends in population changes due to mutations. Cryo-EM consistently detected a smaller population of the RBD-up state than SPR, but the order of mutation effects on population distributions is the same.

The ratio of RBD-down to RBD-up states was used to calculate the Gibbs free energy (ΔG). The lower the ΔG, the higher the RBD-up population. The ΔG was more negative for both dNTD and Delta when compared with that of the WT or dRBD (Figs. 9B & 9C), indicating that the NTD mutations increase the population of the RBD-up conformational state of the spike protein. In contrast, the dRBD ΔG was more positive than the WT, indicating that the RBD mutations decrease the RBD-up conformational states. For the Delta chimeric protein, when both NTD and RBD mutations are present, ΔG is the most negative compared to the other chimeras. This indicates the evolution of NTD mutations as a compensatory mechanism for RBD mutations, which improve receptor binding at the cost of a punitive effect on RBD conformational shifts.

The long-range allosteric communication between the NTD and RBD domains involves significant energetic coupling. The thermodynamic interaction energy between the two domains can be estimated from the double-mutant cycles shown in Figs. 9B and 9C. The nodes represent the relative populations of the RBD-up and RBD-down conformational states as determined from independent experiments on the four chimeric proteins. In both double mutant cycles from SPR and cryo-EM, the sum of free energy changes starting from WT to Delta is the same, irrespective of which off-diagonal direction we travel on the double mutant cycle, which is ΔΔG_WT-dNTD_ + ΔΔG_dNTD-Delta_ = ΔΔG_WT-dRBD_ + ΔΔG_dRBD-Delta_, which represents a perfect double mutant cycle. More importantly, the change in free energy because of both NTD and RBD mutations is greater than the sum of the effect of individual domain mutations (ΔΔG_WT-Delta_ >> ΔΔG_WT-dNTD_ + ΔΔG_WT-dRBD_), which is about -2.32 kJ/mol when measured by SPR (Fig. 9B) and -3.18 kJ/mol when measured by cryo-EM (Fig. 9C), indicating significant energetic coupling underlying the long-range allosteric communication between the two domains.

## Discussion

Over the last two decades, new coronaviruses (CoVs) have consistently emerged, posing a substantial danger to global public health. These include SARS-CoV emerged in 2002, MERS-CoV in 2012, and SARS-CoV-2 in 2019, which resulted in significant infections and deaths.^3,4^ Four other human CoVs are known, typically causing mild to moderate respiratory illness, such as the common cold, with nearly zero fatalities. Viruses evolve through the natural selection of mutations, improving viral fitness on multiple fronts. The spike protein is the most mutated protein in CoVs and is the first point of contact between CoVs and humans. SARS-CoV-2 spike protein has accumulated multiple mutations over the past 5 years, and we can clearly see three mutation hotspots emerging (Fig. 1C). Initial variants have mutations predominantly in the RBD, followed by the NTD, and recent variants show multiple mutations at the S1/S2 interface. We and others have previously shown how mutations in the RBD are evolving, using principles of increased receptor binding and escape from neutralizing antibodies.^10,17,18,20,36,41^ However, very few experimental studies exist on the role of non-RBD mutations, mainly those mutations that are distant from RBD, and how they influence receptor binding and immune escape. Some computational studies suggest long-range allosteric communication between different parts of the spike protein,^31,32^ but these claims have not been tested experimentally. This study probes the role of mutations in the NTD domain, the second most mutated domain of the spike protein, using two independent methods: SPR and cryo-EM, which operate on different physical principles.

SPR probes changes in mass upon binding, whereas cryo-EM provides Coulomb potential maps to visualize structures. Most published SPR studies use 1:1 binding models, which clearly do not explain the biexponential association kinetics of spike proteins binding to ACE2 (Fig. 3). We tested different kinetic schemes based on how spike protein might function, and determined that a two-state conformational change coupled with a binding equilibrium model best describes the SPR data, where the spike protein has to transition from RBD-down to RBD-up conformation before binding to ACE2 (Figs. 4B & 5). The same model also explained the binding kinetics of Class 1, Class 2, and Class 3 antibodies (Figs. 5 & 7). The model also enabled us to calculate the relative populations of the two conformational states of the spike protein (Fig. 9B). Cryo-EM independently confirmed the two conformational states (Fig. 9A), and the populations closely matched those determined by SPR. The populations were then used to construct double mutant cycles (Figs. 9B & 9C) to determine the energetic coupling and long-range allosteric communication between the NTD and RBD domains.

The results presented indicate that NTD mutations affect the conformational dynamics of the RBD in the spike protein. NTD mutations increase the population of RBD-up states of the spike protein, making it more accessible to ACE2 binding as well as to Class 1 neutralizing antibodies that bind only to the RBD-up state (Fig. 5). The main effect is significantly slowing down the kinetics of RBD-up to RBD-down conformational transition with no effect on the RBD-down to RBD-up conformational transition (Fig. 5). The effect of NTD mutations on RBD conformational dynamics is greater in the presence of RBD mutations (Figs. 9B & 9C). Although we observed that the Delta RBD mutations decreased the RBD-up population of the spike protein, the presence of NTD mutations compensated for the decrease. The combined effect of NTD and RBD mutations on RBD conformational dynamics is much greater than that of either domain alone, suggesting long-range allosteric communication between the two domains. We show that this long-range communication involves significant energetic coupling between the two domains (Figs. 9B & 9C). In addition to the RBD conformational dynamics, NTD mutations also increase spike protein expression (Fig. 2C) and stability (Fig. 2I). NTD mutations did not affect the spike protein’s binding to Class 2 and Class 3 neutralizing antibodies, since these antibodies bind in both RBD-up and RBD-down conformations (Fig. 7).

Our results showed that NTD mutations provide the necessary compensation for the energetic consequences of RBD mutations. The development of NTD mutations appears to be an evolutionary mechanism that enables the virus to mitigate the negative effects of certain RBD mutations without compromising viability. This was apparent with the Delta chimera, which showed higher spike protein expression, greater stability, increased receptor binding affinity, and the highest RBD-up population. These findings provide useful insights into the mechanisms underlying the natural selection of mutations that improve viral fitness on multiple fronts. Overall, our findings highlight the synergistic effect of multiple mutations as an essential and adaptive strategy for virus survivability.

## Materials and Methods

### Molecular Cloning of spike chimeras, ACE2, and ScFvs

Prefusion S ectodomain (residues 1-1208) SARS-CoV-2 HexaPro construct^44^ was commercially acquired from Addgene (plasmid #154754) and used in designing spike chimeras. All HexaPro constructs were cloned into a pαH mammalian expression plasmid and contained a C-terminal T4 foldon trimerization motif, HRV3C cleavage site, an 8’ His-Tag, and a Twin-Strep-tag for purification. HexaPro spikes contain the D614G mutation, “GSAS” substituted furin cleavage site (residues 682–685), and six stabilizing proline substitutions F817P, A892P, A899P, A942P, K986P, and V987P. To produce Delta variant NTD (dNTD) chimeras, mutations T19R, T95I, G142D, del156-157, and R158G were added to the wild-type HexaPro using site-directed mutagenesis. Mutations L452R and T478K were similarly added into wild-type and NTD mutant spikes to make the Delta RBD (dRBD) chimera and full Delta NTD-RBD chimera. Non-RBD/NTD mutations were excluded from all chimeras to ensure that any changes in RBD binding or conformation were due to RBD or NTD mutations. Sequences for monomeric hACE2, as well as 3 neutralizing antibodies that span 3 different classes of RBD epitopes, were obtained from Uniprot (hACE2 ID: Q9BYF1) and the Research Collaboratory for Structural Bioinformatics (RCSB) Protein Data Bank (PDB). PDB IDs were used for the following antibodies: Class 1 ScFv LY016 (7C01), Class 2 ScFv Ly555(7KMG), and Class 3 REGN10987 (6XDG). Single-chain variable fragments (ScFv) were designed for each antibody and utilized a triplicate G_4_S linker to connect heavy (V_H_) and light (V_L_) chains. All sequences were codon optimized by Twist Biosciences for expression in mammalian cells. ACE2 and ScFv constructs were cloned into pcDNA3.4-TOPO vectors using BamHI and XhoI restriction sites. Constructs also contained an N-terminal SUMOstar protease fused to a 6x Histidine tag and a human immunoglobulin heavy chain secretory sequence, arranged 5’ to 3’ upstream of the target gene.

### Expression and purification of spike chimeras, ACE2, and ScFvs

Proteins were expressed in Expi293 HEK cells over 5 days. Transfection of spike chimeras was carried out according to the protocols established earlier.^35^ Briefly, cells were seeded at a concentration of 1 x 10^6^ cells/mL in 36 mL Expi293 media, and a transfection mixture containing 4mL OptiPro SFM (Gibco, cat. no. 12309019), 25 μg DNA, and 75 μg polyethyleneimine (PEI) was added dropwise. For ACE2 and ScFvs, cells were seeded at a concentration of 3 x 10^6^ in 30mL and transfected with 30 μg DNA and 90 μg PEI in 1 mL OptiPro SFM. Cultures were incubated on an orbital shaker at 125 rpm in a 37 °C incubator with 8.0% CO_2_. After 5 days, the cultures were centrifuged, and the supernatant was filter-sterilized using a 0.22 μm PVDF filter. Expression levels for the four chimeras were assessed by running culture supernatants on SDS-PAGE, followed by staining with Coomassie Brilliant Blue R-250. Band intensities were quantified using ImageJ and normalized to cell density using Bio-Rad ImageLab. All proteins were purified using Ni-NTA affinity chromatography and eluted in buffer containing 50 mM Tris, 200 mM NaCl, and 150 mM Imidazole, pH 8.0. Spike chimeras were buffer-exchanged into SEC buffer (2 mM Tris, 200 mM NaCl, 0.02% NaN3) and further purified on a Superose-6 10/300 GL SEC column (GE Healthcare, cat. no. 29115046) at a flow rate of 0.5mL/min. For ACE2 and ScFvs, purified protein was buffer exchanged into a buffer containing only 50 mM Tris, 200 mM NaCl, pH 8.0, using a 10,000-30,000 MWCO Amicon Ultra centrifugal filter (Millipore Ltd, cat. no. UFC903024) and incubated at 4 °C overnight with SUMOstar protease to cleave the SUMOstar-bound 6x Histidine tag. Digested proteins were purified again with a Ni-NTA column to remove the cleaved tag. Spike chimeras were buffer exchanged into 1x PBST (10x solution; 80.6 mM sodium phosphate, 19.4 mM potassium phosphate, 27 mM KCl, and 1.37 M NaCl in high purity dH_2_O) pH 7.4, and all other proteins were buffer exchanged into phosphate buffer (50 mM sodium phosphate and 20 mM NaCl) pH 7.4 and stored at -80 ℃. Purity for all proteins was verified using SDS-PAGE.

### CD and Fluorescence Spectroscopy

CD spectra for spike chimeras were obtained using an Applied Photophysics Chirascan Plus spectrometer. A 2 𝜇𝜇M concentration of spike protein in 1x PBS was used for each measurement, and spectra were acquired from 200 nm to 260 nm with a 0.5 mm cuvette. Spectra were recorded at a data interval of 1 nm and an averaging time of 2 seconds. Each spike protein was measured 5 times and averaged to produce the final spectra.

Fluorescence spectra were acquired using a PTI QuantaMaster fluorimeter. A 2 𝜇𝜇M concentration of spike protein in 1x PBS was excited at 280 nm, and the fluorescence emission spectrum was recorded from 300 nm to 400 nm at 1 nm intervals and an averaging time of 1 s/nm. Triplicate spectra were collected and averaged for each spike protein.

### Analytical Ultracentrifugation

The trimeric structure and homogeneity of spike chimeras were confirmed by sedimentation velocity analytical ultracentrifugation (SV-AUC). A 400 µL spike protein solution at a concentration of 0.5 mg/mL was loaded into one sector of a two-sectored cell with a 12 mm centerpiece. The other sector contained 410 µL of 1x PBS, which served as a background reference. Data were collected on a Beckman XL-A ultracentrifuge at 20 C, a wavelength of 280 nm, and a radial step size of 0.003 cm. All cells were run at 40,000 rpm, and a total of 150 scans were collected. The c(s) size distribution analysis was performed using SEDFIT version 16.1 software. Buffer density and viscosity were calculated using SEDNTERP version 3.0.3 software.

### Thermal stability of spike protein variants

The thermostability of wild-type and mutant spike proteins was determined using differential scanning fluorimetry (DSF). A concentration of 1 mg/mL of purified spike in 1xPBS, pH 7.4, was measured in triplicate on a NanoTemper Prometheus Panta (NanoTemper Technologies, Munich, Germany). The intrinsic fluorescence was recorded at 330 and 350 nm while heating the samples from 25 to 90 °C at a rate of 5 °C per minute. The first derivative of the ratio of fluorescence (350/330 nm) as well as the inflection temperatures (*T*_i_) were calculated with Panta Analysis software.

### Surface Plasmon Resonance

A carboxymethyl dextran sensor chip (CMD 200m, Assay Solutions LLC, Boulder, Colorado) was used with a BiOptix 404pi to immobilize twin-strep-tagged spike chimeras for binding experiments. First StrepTactin-XT protein (iba-lifesciences, cat# 2-1204-005) was immobilized onto the reference and binding flow channels using manufacturer recommended conditions.^45^ Next, tagged spike protein (450 kDa) was immobilized onto binding flow channels to a level of ∼300-400 response units (RUs) at a flow rate of 10 μL. To test spike binding affinity, serial dilutions of ACE2 (70 kDa) or ScFvs were injected as the analyte at a flow rate of 50 µL/min. Five analyte concentrations were used for all analytes, with ACE2 concentrations ranging from 250 nM to 7.81 nM and ScFv concentrations ranging from 300 nM to 1.12 nM. For ACE2 binding, the analyte was allowed to flow for 350 seconds to measure association kinetics, followed by a dissociation time of 600 seconds. ScFv binding had an association time of 250 seconds, followed by a 350-second dissociation. All analyte injections for each concentration were repeated in triplicate. For Class 2 and Class 3 ScFvs, binding did not fully dissociate, so the chip surface was regenerated after each analyte injection, per the manufacturer’s recommendations.^45^ After regeneration, the fresh spike was immobilized for the next analyte injection. SPR was also used to generate data for analytical model determination of WT spike binding ACE2 and ScFvs. ACE2 experiments were performed using the same five concentrations as in the binding studies, with two additional concentrations of 100 nM and 165 nM. Class 2 or 3 ScFv experiments used analyte concentrations of 11.11 nM, 33.33 nM, 50 nM, 100 nM, 150 nM, 200 nM, and 250 nM.

Channels with immobilized Strep-Tactin XT alone were used as a reference. All experiments were performed at 25 °C in 1x PBST running buffer, and all proteins were buffer exchanged into fresh buffer prior to analysis. Results for all kinetic rates and affinity are from the average of triplicate experiments. All statistics and data fitting were performed using OriginPro and MATLAB.

### Cryo-EM data collection

To prepare the cryo-EM sample grids, 2 mg/mL purified spikes in PBS pH 7.4 were deposited onto glow-discharged CF-1.2/1.3 carbon-coated copper grids using a Vitrobot Mark IV (Thermo Fisher Scientific) at 100% humidity and 25℃. All grids were prepared with 3 μL protein sample, followed by a wait time of 6 seconds, then blotting of excess liquid with a blot force of -1 and a 3 second blotting time followed by plunge-freezing into liquid ethane. Data for WT and Delta spikes were collected by the Pacific Northwest Center for Cryo-EM (PNCC), and data for dRBD and dNTD were collected by the University of Colorado Anschutz Cryo-EM Core Facility (UCA). PNCC datasets were collected on a 300kV Krios 1 microscope equipped with a Falcon4i camera and a Selectris X energy filter, while UCA data were collected on a 200kV Talos Artica equipped with a Gatan K3 Camera. WT and dRBD data had an accumulated dose of 50 Å^2^/e and calibrated pixel size of 0.73 and 0.91 Å, respectively. For dNTD and Delta, preliminary data collection showed a strong preferred orientation compared to WT and dRBD. To compensate for this, grids were tilted to 30° during collection. The accumulated dose for dNTD and Delta was 50 Å^2^/e with 0.7296 pixel size.

### Cryo-EM 3D reconstruction

All datasets were processed using CryoSPARC v4. A total of 17128, 10413, 6812, and 11180 micrographs were collected for WT, dRBD, dNTD, and Delta, respectively. First, motion correction and contrast transfer function (CTF) estimation were performed, followed by micrograph curation to remove low quality micrographs. Roughly, 10-30% of micrographs were discarded, and the remaining micrographs were used for subsequent processing. Several rounds of particle picking, extraction with a box size of 480 pixels and down sampling of 128 pixels, and 2D classification were performed on the first 1000 micrographs to isolate classes that display clear spike structures. These particles were then used as templates for targeted particle picking, extraction without down-sampling, and 2D classification with the full dataset. Several more rounds of 2D classification were performed to clean up particles, yielding roughly 30-40 classes and between 300,000 and 600,000 particles across all four spike datasets used for ab initio 3D reconstruction. Reconstructed particles were subjected to heterogeneous refinement followed by C3 symmetry expansion and 3D classification using a focused mask over RBD. All 1-RBD up volumes and all 3-RBD down volumes underwent non-uniform refinement and homogeneous refinement, respectively. In cases where volumes showed missing resolution in RBD, a local refinement was run with a focused mask over RBD, with the fulcrum coordinates set at the base of the mask. Finally, local resolution estimation was run on all final volumes. 3D variability was run on the final 1RBD up volumes to confirm that the population was homogeneous.

## Acknowledgements

We thank Francisco Asturias & Shaodong Dai, University of Colorado Anschutz, Vignesh Kasinath, University of Colorado Boulder, and Jason McLellan, University of Texas at Austin, for many helpful discussions on cryo-EM methods and data analysis. We thank David Bain and Carlos Catalano at the University of Colorado Anschutz for many discussions on the analysis of SPR kinetic data. We thank Arthur Goldsipe of MathWorks for his guidance on numerically fitting the SPR association and dissociation kinetics to complex kinetic schemes using compartmental analysis in MATLAB. We thank Vaibhav Upadhyay and Casey Patrick from our lab for many discussions on the structure and function of the coronavirus spike proteins. We thank Walter Englander, University of Pennsylvania, for his critical reading of the manuscript.

This work was funded by the Associate Dean for Research, Skaggs School of Pharmacy and Pharmaceutical Sciences, University of Colorado Anschutz, and USA National Institutes of Health (NIH) grant R21 5R21AI183195. The Pacific Northwest Cryo-EM Center was funded by the NIH grant P2C-GM148560, and the cryo-EM Structural Biology Shared Resource Facility, University of Colorado Anschutz, was funded by the NIH grant P30CA046934.

